# ParA and its functions that go beyond chromosome segregation in *Caulobacter crescentus*

**DOI:** 10.1101/2022.05.18.492544

**Authors:** Inoka P. Menikpurage, Stephanie G. Puentes-Rodriguez, Rawan A. Elaksher, Paola E. Mera

## Abstract

Maintaining the integrity of the chromosome after the completion of each cell cycle is paramount for bacterial survival. Mechanistic details remain incomplete for how bacteria manage to retain intact chromosomes in each daughter cell after each cell division. In this study, we examined the partitioning protein ParA and its functions on chromosomal maintenance that go beyond triggering the onset of chromosome segregation. Our data demonstrate that ParA can promote the onset of chromosome replication initiation in Caulobacter crescentus cells with sub-physiological levels of the replication initiator protein DnaA. Increasing the cellular levels of ParA results in over-initiation of chromosome replication in this bacterium. We show that the ability of ParA to impact replication initiation is independent from ParA’s ability to trigger the onset of chromosome segregation. Surprisingly, our work revealed that perturbing the balance of the components of ParA’s nucleotide-dependent cycle can have severe defects in cell cycle coordination and can potentially be lethal to the cell. Increasing the levels of different forms of ParA also impacted cell length independent of their replication initiation frequencies. Our results, together with past observations, suggest a model where ParA can serve as a checkpoint coordinating various cell cycle events involved in maintenance of the chromosome.

## INTRODUCTION

The survival of bacteria relies on their ability to preserve the integrity of their genome. Maintaining intact copies of the chromosome after each cell division requires exquisite temporal and spatial coordination of different cell cycle events. In bacteria, this coordination must be accomplished while the chromosome is being replicated and segregated concurrently [1]. Any mistake in temporally coordinating chromosome replication, segregation, and cytokinesis can result in cell death. Although details of each of these processes have been elegantly revealed, mechanistic details of how these events are coordinated, temporally and spatially, with each other remain limited. In this study, we focused on the coordination between the onset of chromosome replication and the onset of chromosome segregation.

The onset of chromosome replication – a major regulatory step in the cell cycle – is catalyzed by the conserved AAA+ ATPase replication initiator DnaA [2]. DnaA-ATP oligomerizes at the origin of replication (*ori*) and opens the double stranded DNA helix allowing for the replication machinery to assemble and replicate the chromosome bidirectionally [3]. The ability of DnaA to initiate chromosome replication is regulated at a multitude of levels that include autoregulation, deactivation, sequestration, titration, and proteolysis [4]. In addition to catalyzing this essential step for DNA synthesis, DnaA and/or replication initiation have been shown to influence other key cellular events including chromosome segregation, cell cycle progression, cytokinesis, cell size, and gene expression [5]. These various connections expose the complexity of the highly interconnected network that links one of the first committed events in this developmental process, the onset of chromosome replication, with the rest of the cell cycle. However, mechanistic details at the molecular level of how the initiation of chromosome replication links to various cell cycle events have remained limited.

The partitioning system ParABS is involved in the onset of chromosome segregation. The ParABS system was first discovered and characterized in the segregation and maintenance of low copy plasmids [6; 7; 8]. Most bacterial species (>70 %) encode the parABS system [9]. ParABS is composed of three parts: a centromere-like chromosomal locus *parS* and two DNA-binding proteins ParA (ATPase) and ParB (CTPase). ParA bound to ATP forms dimers that bind DNA nonspecifically and are released from DNA through the interaction with ParB [10; 11]. Different lines of evidence have demonstrated connections between the onset of replication and segregation [1]. For instance, the chromosomal locus parS is commonly found near the chromosomal origin of replication suggesting a potential proximity link between the onset of replication and segregation [7; 9; 12]. In *Bacillus subtilis* and *Vibrio cholera*, the partitioning ParA protein has been shown to regulate the onset of chromosome replication [13; 14]. In *B. subtilis*, a variant of ParA (known as Soj) that remains a monomer was shown to directly interact with DnaA and inhibit its ability to oligomerize at *ori* and thus prevent the onset of chromosome replication [15]. Furthermore, the stability of DnaA bound at *ori* increases in *B. subtilis* with *soj* (*parA*) knocked out resulting in over-initiation of replication [16].

In *C. crescentus* the ParABS system is essential for chromosome segregation and thus essential for viability [17]. The asymmetric cell cycle of *C. crescentus* has facilitated extensive work on the ParABS revealing fine details of the connection between the onset of replication and segregation. For instance, once replication initiates at *ori*, chromosome segregation is not triggered until the *parS* chromosomal locus is replicated [18] (Figure 1). The relatively high concentrations of ParB molecules bound at *parS* are necessary to trigger the ATPase activity of ParA [19]. Dimers of ParA interact with the chromosome nonspecifically, forming a cellular gradient that retracts as one copy of the replicated *parS* (decorated with ParB) segregates to the opposite pole [20; 21; 22] (Figure 1). Connections between the onset of replication and segregation have been proposed in *C. crescentus*. For instance, DnaA, aside from binding *ori*, was shown to bind *parS* directly and promote the onset of chromosome segregation in a ParA-dependent manner [23]. Interestingly, the nucleoid-associated protein GapR, similarly to DnaA, binds at *ori* and *parS* coordinating the onset of chromosome replication with the asymmetric chromosome separation [24]. Although *C. crescentus* serves as a powerful system to investigate the coordination between the onset of chromosome replication and the onset of segregation, the potential role of ParA as a modulator of chromosome replication initiation has not yet been reported.

**Figure 1:**
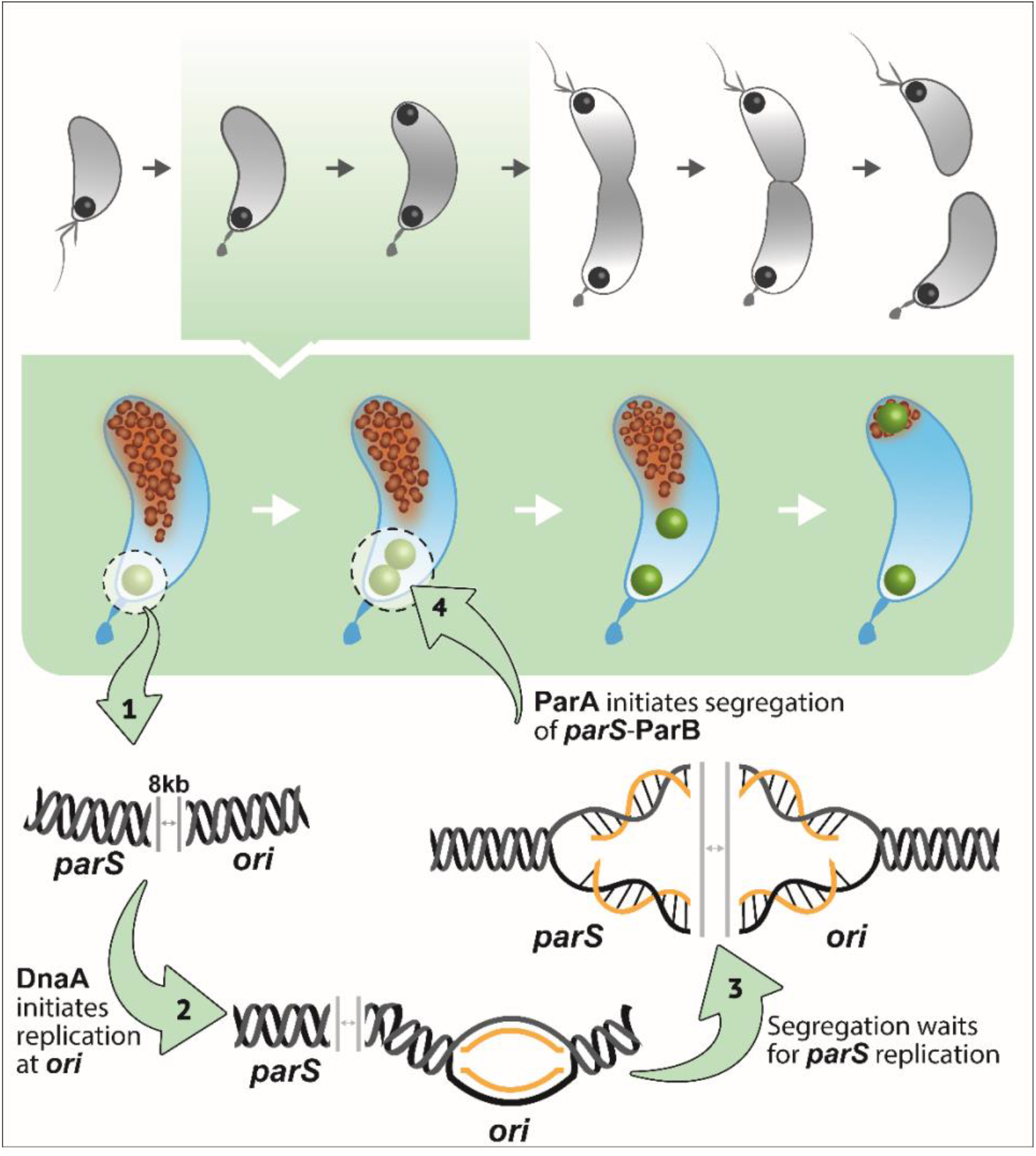
Schematic of *Caulobacter crescentus* asymmetric cell cycle, highlighting the dynamics of the onset of chromosome replication and segregation. Nondividing cells contain the origin of replication (ori) and the centromere-like region (parS) (black dot) near the stalked pole (top panel). The origin of replication and the ParB/*parS* complex segregate first towards the new pole in a ParA-dependent fashion, following the cloud-like gradient of ParA retract (middle panel). DnaA initiates chromosome replication at ori. ParA does not initiate chromosome segregation until parS is replicated (bottom panel).

In this study, we discovered that ParA in *C. crescentus* can also promote the onset of replication, albeit not necessarily through direct interactions with DnaA. Our data reveals that ParA can promote the onset of replication at sub-optimal levels of DnaA. We demonstrate that ParA levels can significantly alter the frequency of replication initiation in a DnaA-ATP dependent manner. We used a set of ParA variants to demonstrate that ParA can promote replication initiation independent of ParA’s ability to interact with DNA or hydrolyze ATP. Thus, ParA can promote the onset of replication independently of ParA’s ability to trigger the onset of chromosome segregation. Remarkably, we discovered that potential expression of the monomer apo-ParA can result in lethal outcomes. Collectively, these data provide new insights into the complex coordination between these two fundamental events involved in the maintenance of chromosome integrity.

## RESULTS

### ParA perturbs frequencies of chromosome replication initiation

*Caulobacter* initiates chromosome replication only once per cell cycle, regardless of the type of growth media (minimal vs. rich). We previously reported an intriguing observation where cells with high levels of the partitioning ParA protein were able to initiate chromosome replication in cells containing insufficient levels of the replication initiator DnaA [25]. This observation suggested a potential link between chromosome replication and chromosome segregation. To further analyze that observation, we examined the frequency of replication initiation in cells overexpressing *parA*. To do that, we constructed a merodiploid *Caulobacter* strain expressing *parA* from the chromosomal xylose inducible promoter in addition to the native copy of *parA* under its own promoter (PM541: NA1000, *xylX::parA, parB::cfp-parB*). To track the initiation of chromosome replication, we fluorescently labeled the partitioning protein ParB (CFP-ParB) that binds the centromere-like region *parS* [26]. Because the *parS* locus is found near the origin of replication [18], we can infer the frequency of replication initiation by counting the number of CFP-ParB foci per cell [26]. Our data revealed that cells (PM541) overexpressing *parA* display multiple CFP-ParB foci, suggesting that these cells were over-initiating chromosome replication potentially due to increased levels of ParA (Figure 2A). The addition of the same xylose concentrations to a strain with an empty-vector control (PM566: NA1000, *xylX*::empty-vector, *parB::cfp-parB*) resulted in cells behaving like the wild-type control displaying only one or a maximum of two CFP-ParB foci per cell (Figure 2B).

**Figure 2:**
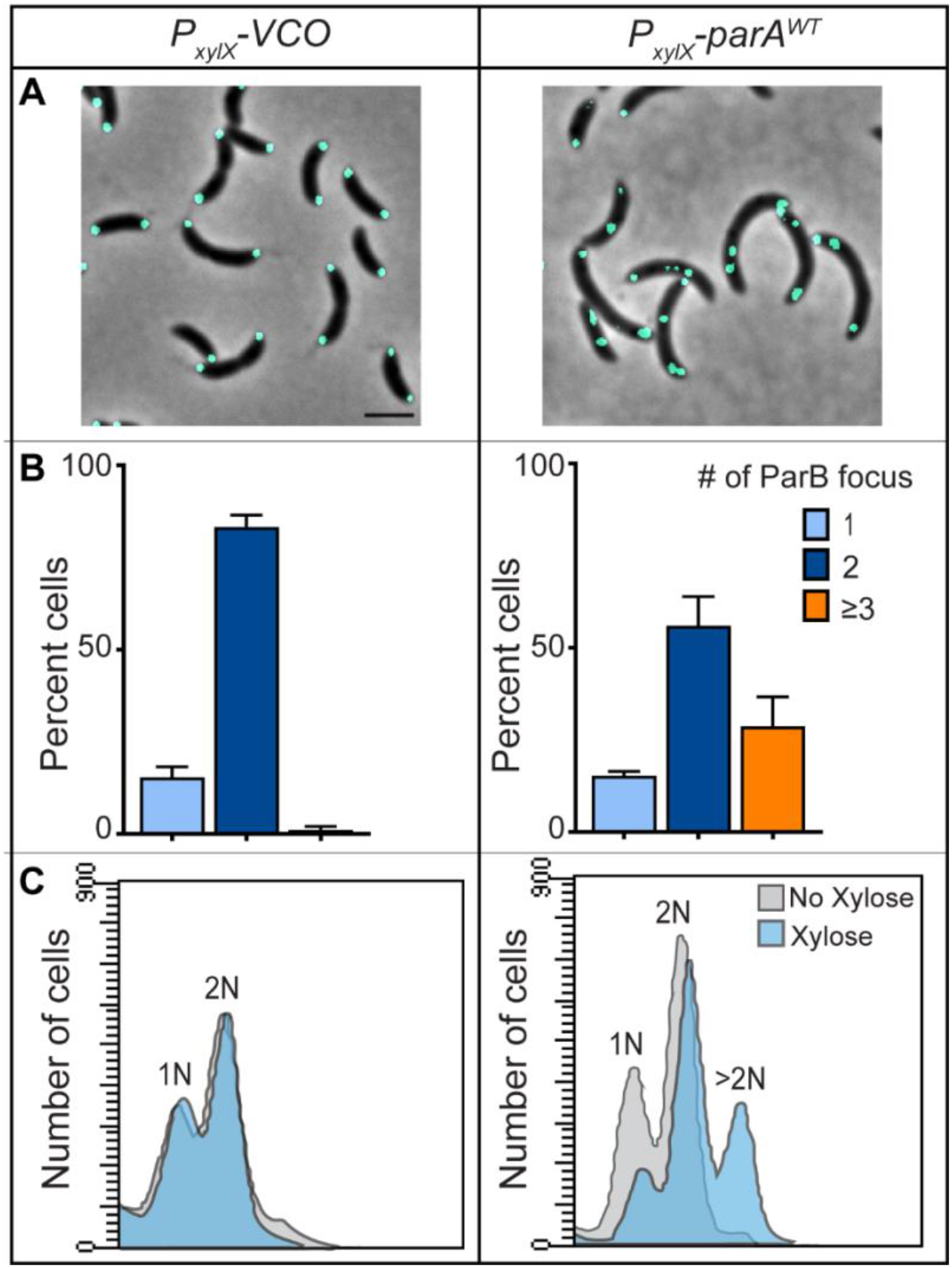
High levels of ParA^WT^ promote over-initiation of chromosome replication in *C. crescentus*. Isolated swarmer cells of CB15N, *parB::CFP-parB* cells with empty vector control, and *xylX::ParA*^*WT*^ in M2G (2ml, OD_600_ ∼ 0.1) medium were supplemented with 0.1% xylose at t = 0 h and incubated at 30°C in a roller-shaker. Phase-contrast fluorescence micrographs were obtained at t = 3 h and blind-quantified the number of CFP-ParB foci per cell which indicates the number of *ori*s. (A) Micrograph of CB15N, *parB::CFP-parB* cells with empty vector control, and *xylX::ParA*^*WT*^ expressing ParA^WT^ for 3h. (B) Bar graphs of the percent number of ParB foci (*ori*s) per cell. (C) Flow cytometry profiles of cells after 3 h incubation. Micrograph scale bar – 2μm. Bar graph data are from three independent experiments with error bars of mean ± standard deviation (SD).

We used two additional methods to determine the impact of *parA* overexpression on replication initiation. Firstly, we quantified the number of *ori* independent of ParB by testing a *Caulobacter* strain in which the area near the origin of replication is fluorescently labeled with TetR-eYFP and a *tetO* array [27]. Consistent with our ParB-labeled cells, increasing the levels of *parA* expression in this *ori* labeled strain resulted in multiple *ori* foci (Supplemental Figure 1). Secondly, we analyzed the chromosomal content of cells overexpressing *parA* using flow cytometry. Cells that were induced for *parA* overexpression displayed an additional curve of >2 chromosomes per cell (Figure 2C). This increase in chromosomal number was not observed in cells with vector-control only (VCO) treated under the same conditions. These cumulative data confirmed that increasing levels of *parA* expression results in cells comprising multiple origins of replication.

Furthermore, we determined the levels of ParA that were able to promote the over-initiation of chromosome replication using our *parA* merodiploid strain. To track the over-expression levels of *parA*, the protein ParA was tagged with the peptide M2 (DYKDDDDK) at the C-terminus. We first confirmed that the expression of ParA-M2 from *parA*’s native promoter had no effects on growth (Supplementary Figure 2A). Cells expressing parA-M2 from the xylose promoter retain their ability to trigger over-initiation of chromosome replication (Supplementary Figure 2B). To determine the levels of ParA, we compared native parA-M2 levels versus xylX::parA-M2 using Western blots. Our data revealed that overexpression of *parA* from the xylose promoter results in a 5-fold increase of ParA levels after 3 h of induction compared to VCO (Supplementary Figure 2CD).

### DnaA-ATP is required for ParA’s impact on replication initiation

Given that increased levels of ParA result in over-initiation of replication, we tested the possibility of ParA promoting the onset of chromosome replication independently of the canonical replication initiator DnaA. Because DnaA is essential for viability, we tested this DnaA-independent replication hypothesis using a strain where the levels of DnaA can be depleted. This depletion is accomplished by replacing the native *dnaA* copy with a streptomycin cassette and incorporating a *dnaA* copy downstream of the inducible chromosomal promoter vanillate (PM109) [23; 28]. We engineered into PM109 a second copy of *parA* under the inducible promoter xylose (PM542: *dnaA::spec, vanA::dnaA, xylX::parA, parB::cfp-parB*). These cells (PM542) were depleted of DnaA for 3 h and then induced for *parA* overexpression for an additional hour. Cells depleted of DnaA displayed only one CFP-ParB focus, consistent with their inability to initiate chromosome replication in the absence of DnaA (Figure 3A). In our positive control condition, ∼98% of cells that had DnaA depleted (3h) and then repleted (1h) displayed 2 CFP-ParB foci showing that DnaA has no problem initiating replication after the depletion period. In the potential scenario where ParA can initiate chromosome replication independently of DnaA, PM542 cells depleted of DnaA and then exposed to xylose to induce expression of *parA* would also display two CFP-ParB foci per cell. However, our data revealed that cells overexpressing *parA* (+ xylose) in the absence of DnaA (no vanillate) display only one CFP-ParB focus per cell (Figure 3A). High levels of ParA had no effect on the number of CFP-ParB foci in cells that lacked DnaA. These data indicate that ParA cannot initiate chromosome replication and its apparent role on the onset of chromosome replication is DnaA-dependent.

**Figure 3:**
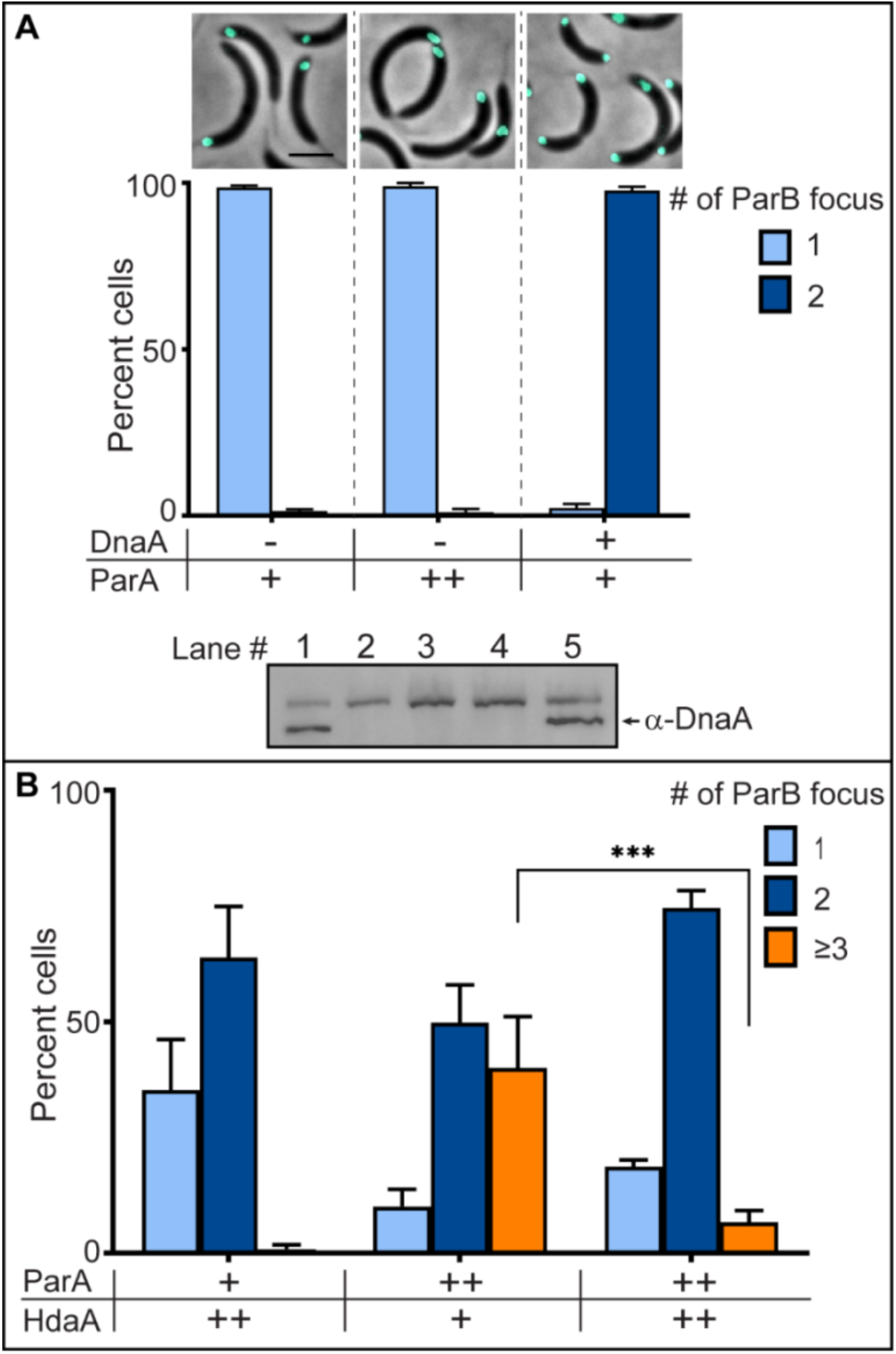
DnaA-ATP is required for ParA to promote replication initiation. Swarmer cells of *C. crescentus* CB15N, *parB*:*cfp*-*parB*, Δ*vanA, dnaA*::Ω, *vanA*::*dnaA*,and *xylX::parA*^*WT*^ (merodiploid ParA cells) were isolated from a culture grown in M2G with vanillate 250 μM (40 ml, OD_600_ ∼ 0.3). Swarmer cells resuspended in M2G were split into 3 (2 ml each) and incubated at 30°C in a roller-shaker for 3 h to deplete DnaA. Upon DnaA depletion, tubes were incubated with no inducer, xylose (0.1%), and vanillate (250 μM) for 1 h, and Phase-contrast fluorescence micrographs were obtained. (A) Micrographs and quantification plots of the number of CFP-ParB foci/*ori* per cell: depleted for 4h (left), cells expressing ParA^WT^ under xylose promoter (middle) and expressing DnaA (right) showing only cells expressing DnaA initiate the replication (2 CFP-ParB foci). Western blot (bottom) of DnaA levels of each cell sample: equal amount of cells loaded onto lane 1 – 5: swarmer cells, DnaA depleted cells for 3 h, DnaA depleted cells for 4 h, cells expressing ParA^WT^ for 1 h, and cells expressing DnaA for 1 h. (B) Bar graphs of CFP-ParB foci quantification of merodiploid ParA and HdaA cells (CB15N, *parB*:*cfp*-*parB*, Δ*vanA, vanA*::hda*A, xylX::parA*^*WT*^) cells expressing HdaA and/or ParA. Cells were grown in M2G (40 ml) up to OD_600_ ∼ 0.3. Vanillate (250 μM) was added to induce HdaA and incubated at 30°C for 1h before synchrony. Swarmer cells resuspended in M2G were split into 3 tubes (2 ml per tube) and the cultures were supplemented with vanillate (250 μM), the second tube with xylose (0.1%) and the third tube with both vanillate (250 μM) and xylose (0.1%) and incubated for 3 h. Fluorescence micrographs obtained at 3 h were used to quantify the number of CFP-ParB foci. Expression of HdaA reduced the frequency of over-initiation suggesting DnaA-ATP is required for ParA’s role in replication initiation. Micrograph scale bar – 2 μm, (+) Wild-type gene expression, (++) Induced protein levels under wild-type background. Bar graph data are from three independent experiments with error bars of mean ± standard deviation (SD).

We examined another possible scenario that would explain the over-initiation of chromosome replication where ParA stimulates replication initiation with deactivated DnaA (DnaA-ADP). To test this hypothesis, we quantified the frequency of replication initiation in cells that contained predominantly high levels of DnaA-ADP compared to DnaA-ATP. To increase the ratio of DnaA-ADP/DnaA-ATP, we increased the levels of the protein that promotes the ATPase activity of DnaA, HdaA [29]. We constructed a merodiploid *hdaA* strain with the additional copy of *hdaA* under the vanillate inducible promoter (PM607: *vanA::hdaA, xylX::parA, parB::CFP-ParB*). PM607 cells were pre-induced for overexpression of *hdaA* for 1h. Post *hdaA* induction, the frequency of replication initiation was quantified. In our control treatment, cells exposed to continuous overexpression of *hdaA*, but no *parA*, displayed either one or two *ori*s (Figure 3B), consistent with what has previously been reported [29; 30]. Cells overexpressing *parA* alone displayed over-initiation of replication in ∼40 % of cells revealing that cells can recover their levels of active DnaA within one hour. However, when the expression of *hdaA* was co-induced with *parA*, the rates of over-initiation of replication dropped (Figure 3B). These data revealed that when the levels of DnaA-ADP are the predominant form of DnaA in the cell, ParA is unable to promote the over-initiation of chromosome replication suggesting that ParA has no role in the activation-deactivation dynamics of DnaA.

### ParA promotes the initiation of replication at sub-optimal levels of DnaA

We analyzed the potential ability of ParA to promote chromosome replication initiation in cells expressing sub-optimal levels of DnaA. First, we determined the minimum induction levels necessary for DnaA to initiate chromosome replication by depleting DnaA levels first and then titrating levels of vanillate (van) to induce different levels of *dnaA* expression (Supplemental Figure 3). The concentration of vanillate typically used to induce expression of the vanillate promoter is 250 μM [31]. Within 1 h of induction (post-depletion), cells expressing *dnaA* from the Pvan promoter at 10 μM vanillate displayed the same frequency of replication initiation as 250 μM van treatment. Vanillate concentrations below 0.5 μM were insufficient to trigger replication initiation after 1 h induction (Supplemental Figure 3). Using this concentration (0.5 μM van) of inducer that was insufficient to allow replication initiation, we tested whether high levels of ParA could influence the ability of these cells to initiate replication. Our xylose-VCO strain depleted of DnaA and then exposed to xylose displayed only one ParB focus per cell demonstrating that the addition of xylose has no impact on the onset of chromosome replication under the conditions tested (Figure 4A). Notably, when cells contain sub-optimal levels of DnaA and high levels of ParA, the frequency of chromosome replication initiation increases as evidenced by the higher number of cells displaying two CFP-ParB foci. These data are consistent with our previous observation that *parA* overexpression results in cells initiating replication at sub-optimal DnaA levels [25].

**Figure 4:**
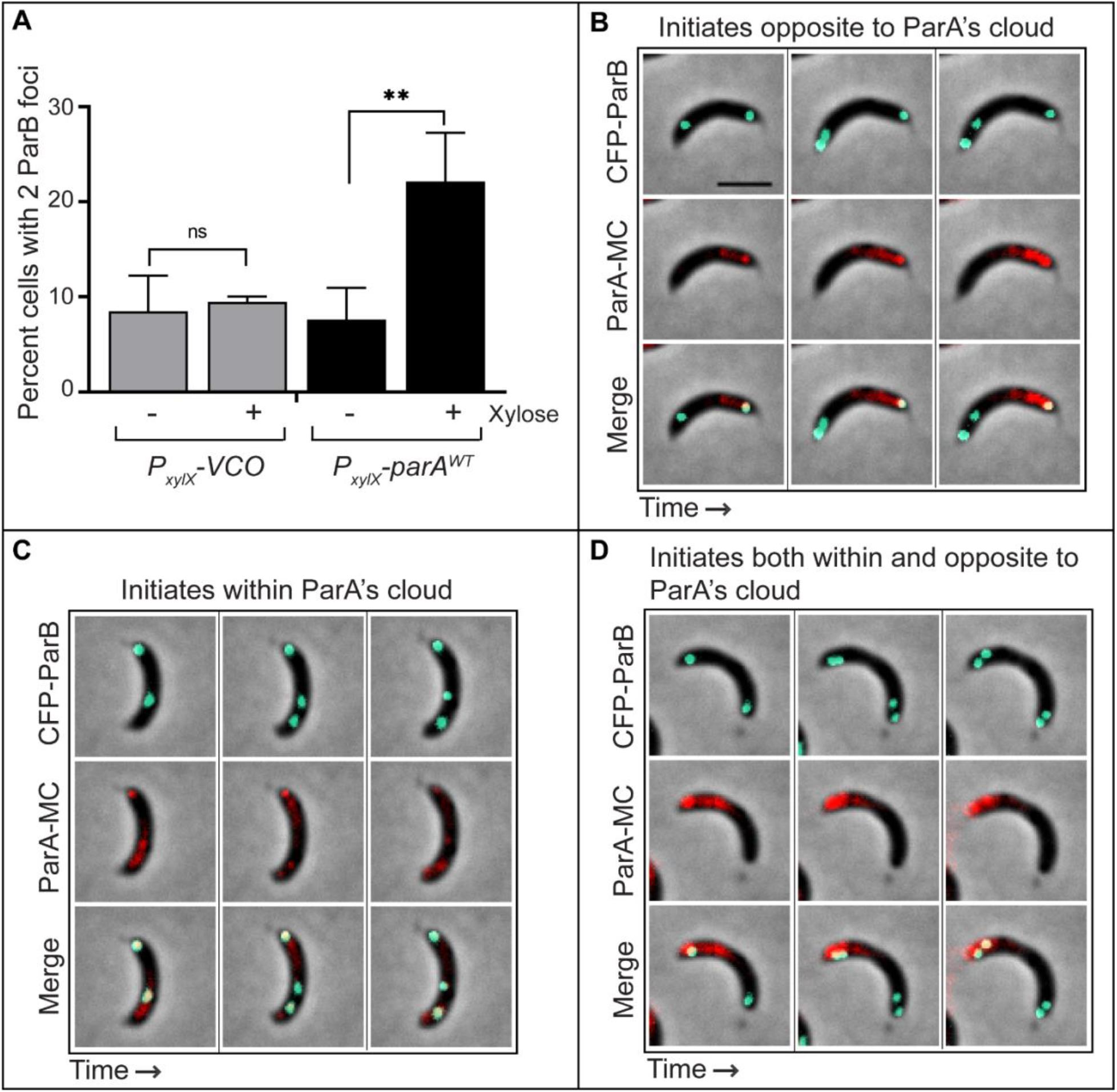
ParA’s impact on replication initiation. (A) Plots of number of cells with 2 CFP-ParB foci/*ori* of CB15N, *parB*:*cfp*-*parB*, Δ*vanA, dnaA*::Ω, *vanA*::*dnaA, xylX::empty-vector* and *xylX::ParA*^*WT*^ expressing ParA^WT^ for 3h with vanillate (0.5 μM) and +/- xylose (0.1 %). Bar graph data are from three independent experiments with error bars of mean ± standard deviation (SD). (B) Phase-contrast fluorescence micrographs of CB15N *parB*:*cfp*-*parB, xylX::parA-mCherry* (MC) cells expressing ParA^WT^-MC. Swarmer cells were incubated with xylose (0.1 %) for 1.15 h and spotted (t= 0 h) on M2G agar pad containing xylose (0.1 %). Time-lapse micrographs were obtained every 15 min over 2.30 h. Cells in (B) showing the initiation of replication opposite to, (C) within and (D) both within and opposite to ParA’s cloud. Time points: (B) 30, 60, and 90 min, (C and D) 45, 75, and 105 min. Micrograps data are representative of three independent experiments. All cell cultures grown up to OD_600_ 0.3 were supplemented with xylose (0.1 %) prior to synchrony to separate swarmer cells.

To investigate a potential mechanism that requires direct protein-protein interactions, we analyzed the localization of ParA and DnaA in cells that over-initiated chromosome replication. We fluorescently labeled ParA (ParA-mCherry) and analyzed its localization patterns. We determined where over-initiation occurs (where *ori-parS* localizes) inside the cell in reference to ParA’s gradient. To do this, we performed a timelapse to track before and after the over-initiation of replication. Our data revealed that over-initiation of replication occurred independently of the location of ParA. We observed the second onset of replication to be triggered from *oris* located away from ParA’s cloud, within ParA’s cloud, or within the same cell away and at ParA’s cloud (Figure 4BCD). These data suggest that ParA’s effect on replication initiation is not likely due to a direct DnaA-ParA interaction in *Caulobacter*.

### Changes in frequency of replication initiation result in unaltered growth rate

*E. coli* is known to initiate multiple rounds of replication when grown in rich media that allows it to grow and divide faster than the time that is necessary to complete the replication of its chromosome [32]. Given that our cells were initiating multiple rounds of replication per cell cycle, we tested whether *Caulobacter* is able to double in shorter times in cells overexpressing *parA*. Using growth curves, our data revealed that the increased frequency of replication initiation had no impact on the doubling time compared to our wild-type control (Figure 5A). We next tested whether rich media could impact the doubling time in cells overexpressing *parA*. We first confirmed that increased levels of ParA cause over-initiation of replication in rich media (Supplemental Figure 4). Our growth curves revealed that the over-initiation of replication showed no difference in doubling time when grown in minimal or rich media. Furthermore, we tested whether the overall progression of the cell cycle was altered when ParA levels were increased by using *Caulobacter’s* master cell cycle regulator CtrA as the indicator. The levels of CtrA are tightly regulated at multiple levels over the cell cycle [33]. Our data revealed that the cell cycle profile of CtrA levels remained the same in cells overexpressing *parA* just like the wild-type control (Figure 5B). These data suggest that high levels of ParA do not disturb the CtrA checkpoint of cell cycle progression. These data are consistent with previous analyses where changes in levels of the par system resulted in no changes in the localization of CtrA’s regulator (CckA) nor changes in expression levels of genes regulated by CtrA [34]. Furthermore, our results suggest that the ability of *E. coli* to increase their growth rates in rich media requires potentially the coordination of multiple events in the cell cycle beyond the over-initiation of chromosome replication.

**Figure 5:**
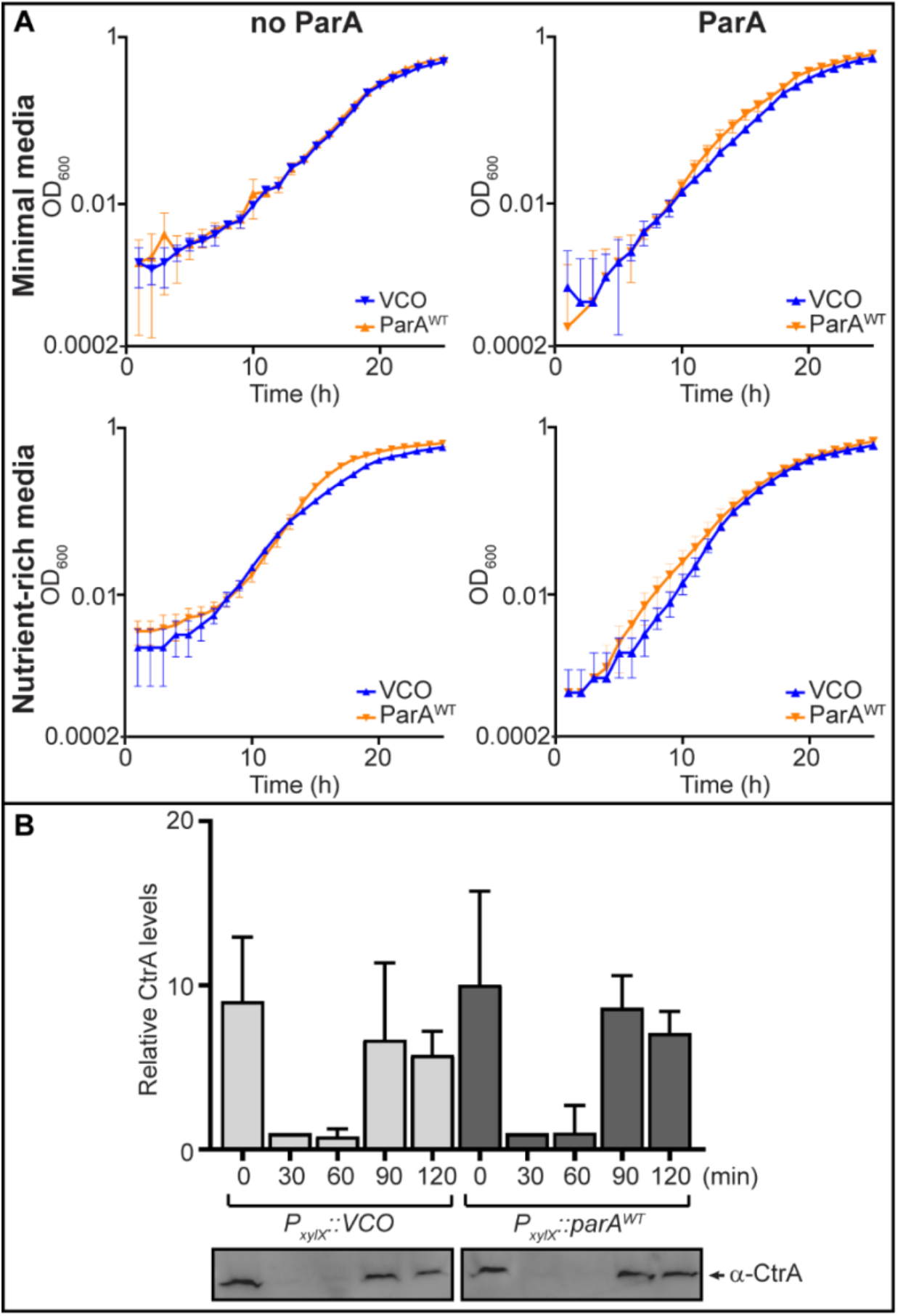
High levels of ParB do not disturb growth. (A) Growth curves of *C. crescentus* CB15N, *parB::CFP-parB* cells with empty vector control, and *xylX::ParA*^*WT*^ in minimal (M2G) and rich (PYE) media in the absence or presence of 0.1 % xylose (expressing ParA^WT^). (B) Bar graph of relative levels of CtrA expression in cells with VCO and higher levels of ParA^WT^ in M2G media supplemented with xylose (0.1%). Cells grown in M2G media were supplemented with xylose (0.1%) 1 h prior to synchrony to induce the expression of ParA. Swarmer cells were resuspended in M2G supplemented with xylose (0.1 %) and incubated at 30°C in a roller-shaker for 2 h. Cell pellets (normalized to OD_600_ 0.2) were saved at t = 0, 30, 60, 90 and 120 min for western blots. Data are from three independent experiments with error bars of mean ± standard deviation (SD).

### ParA levels are linked to the initiation of replication and cell length regulation

We next investigated the potential connection between replication initiation and cell length. In *E. coli*, the over-initiation of replication results in longer cells [35]. However, details of this connection as to which event (over-initiation of replication vs. cell length) causes which have remained elusive in the field. Our data revealed that increasing levels of ParA, which results in over-initiation of replication, also increases cell length (Figure 6A). One potential scenario is that high ParA levels cause the over-initiation of replication, which then cause cells to get longer. Further analyzes of cell size presented below provide more insights into this connection between ParA and cell length.

**Figure 6:**
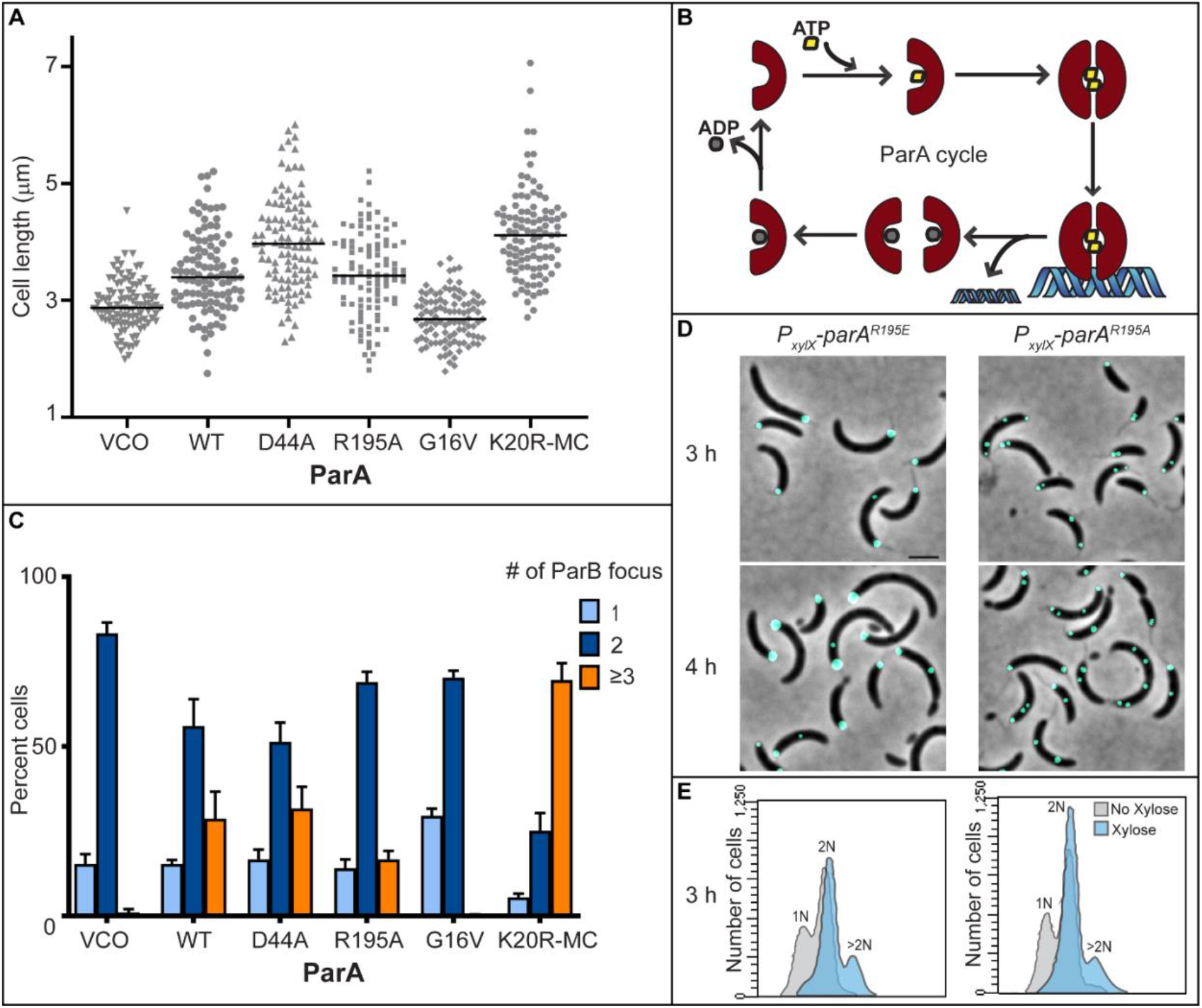
ParA levels are linked to initiation of replication and cell length regulation. (A) Scatter dot lot of cell length of ParA variants in the presence of xylose (0.1 %) for 3 h. Mix population of CB15N *parB cfp*-*parB* with empty vector control (VCO), *xylX::parA*^*WT*^, *xylX::parA*^*D44A*^, *xylX::parA*^*R195A*^, *xylX::parA*^*G16v*^, and *xylX::parA*^*K20R*^-MC was incubated at 30°C for 3 h in M2G media with xylose (0.1 %) before obtaining micrographs. Cell length was analyzed using MicrobeJ software. (B) Cartoon representation of the ParA cycle. (C) Quantification plots of percent cells of CFP-ParB foci in cells expressing ParA variants. Swarmer cells were incubated with xylose (0.1 %) in M2G media. Bar graph data are from three independent experiments with error bars of mean ± standard deviation (SD). (D) Micrographs of CB15N *parB*:*cfp*-*parB, xylX::parA*^*R195E*^ and *xylX::parA*^*R195A*^ cells incubated for 3 and 4 h in M2G + xylose (0.1 %). (E) Flow cytometry profiles of cells after 3 h incubation. Micrograph scale bar – 2μm.

### ParA’s impact on replication initiation and cell length is independent of chromosome segregation

To further characterize this potential new role of ParA, we focused on ParA’s protein dynamics (Figure 6B) by analyzing a set of ParA variants that are unable to trigger chromosome segregation. Given that ParA is essential for chromosome segregation [17; 18], the expression of each ParA variant was regulated by the chromosomal xylose promoter in an otherwise wild-type background. These ParA variants have been previously shown to express stable proteins in *Caulobacter* [20; 36]. To determine whether ParA can promote replication initiation independent of its ability to promote chromosome segregation, we analyzed the variant ParA^D44A^. This ParA variant can bind ATP and dimerize, but it is unable to hydrolyze ATP [13; 20; 37]. In other words, ParA^D44A^ is trapped as ParA dimers bound to ATP. In *Caulobacter*, the ATPase activity of ParA is required for chromosome segregation [18]. Expressing this dimer variant ParA^D44A^ from the xylose promoter resulted in cells with more than two CFP-ParB foci per cell (Figure 6C), similar over-initiation frequencies as those observed with the over-expression of ParA^WT^ (wild-type ParA). Notably, the length of cells expressing ParA^D44A^ was longer than cells overexpressing ParA^WT^, even though both displayed similar frequencies of over-initiation of replication (Figure 6A). These data suggest that the over-initiation of replication is not necessarily directly correlated only to cell length. Overall, these data revealed that ParA is proficient in promoting the over-initiation of replication, independent of ParA’s ability to trigger chromosome segregation.

We also examined whether ParA’s ability to bind DNA is required for ParA’s role in the regulation of replication initiation and cell length. Dimers of ParA bound to ATP interact with DNA nonspecifically, which is required for ParA’s ability to trigger chromosome segregation in *Caulobacter* [17; 18]. We focused on the conserved residue Arg195, previously shown to be critical for ParA’s ability to bind DNA [36; 38]. ParA^R195E^ can bind ATP and dimerize, but it is unable to interact with DNA [36]. Our data revealed that cells expressing ParA^R195E^ displayed only one CFP-ParB focus, initially suggesting that ParA^R195E^ was unable to promote replication initiation (Figure 6D). Over time, however, the single focus we observed with cells expressing ParA^R195E^ became bigger and brighter. To further characterize these abnormally large CFP-ParB foci, we quantified the chromosomal content of cells expressing ParA^R195E^. Our flow cytometry analyses revealed that cells expressing *parA-(R195E)* contained more than two chromosomes (Figure 6E). Thus, cells displaying the larger CFP-ParB focus overtime can indeed over-initiate chromosome replication but are presumably unable to separate the two *oris* from each other. To further confirm these findings, we analyzed another variant unable to bind DNA with an Ala modification at the same R195 residue. Consistent with our FACS findings, cells expressing *parA- (R195A)* displayed multiple CFP-ParB foci similar to cells overexpression the wild-type *parA*. Further characterization of these two ParA variants (R195E and R195A) is necessary to determine the mechanism that explains the differences in their ability to separate the multiple *ori/parS* regions. Collectively, we can conclude that the ability of ParA to hydrolyze ATP and/or to bind DNA are not necessary for ParA’s effect on the onset of chromosome replication and cell length. These data suggest that ParA itself, and not necessarily its ability to trigger chromosome segregation, indirectly impacts replication initiation.

### The apo-ParA monomer severely disrupts the progression of the cell cycle

Given that chromosome segregation activity was not required for ParA’s impact on replication initiation, we examined the effect of monomers of ParA. We focused on two forms of ParA monomers: monomer ParA bound to ATP and monomer apo-ParA. To determine whether the monomer ParA-ATP is involved in promoting replication initiation, we analyzed cells expressing the variant encoding ParA^G16V^ and quantified the frequencies of over-initiation of chromosome replication. The glycine to valine change (G16V) renders this ParA variant able to bind ATP but unable to form dimers due to steric clash [37]. Unlike the over-expression of wild-type ParA, cells expressing *parA-G16V* did not over-initiate replication nor did it impact cell length. Cells expressing *parA-G16V* displayed the same frequencies of replication initiation as cells with empty vector control: only 1 or a maximum of 2 CFP-ParB foci per cell (Figure 6C). These data suggest that monomers of ParA-ATP cannot alter the onset of chromosome replication and/or deregulation of cell length.

Our analyses of the monomer apo-ParA revealed surprising results. To examine the apo-ParA, we focused on the conserved Arg20 residue located in the Walker A motif critical for nucleotide binding [37; 39]. We first noticed the potential dominant lethal effect of this variant while trying to construct the *parA* merodiploid strain with xylose-controlled *parA(K20R)* expression. After multiple attempts, we were unable to get any transformants of *Caulobacter* cells with *parA-K20R*. Our transformant plates either had no colonies or sometimes plates contained one or two colonies that upon sequencing revealed the K20R mutation reverted to wild-type *parA* (Figure 7A). We also tried the construction of this strain using minimal and rich media and at different temperatures in hopes that slower growth would allow cells to retain the *parA(K20R)* construct. However, all these attempts were unsuccessful. We performed this transformation more than seven times but quantified the number of transformants only in the last two trials. We were able to circumvent this potential lethality by adding a mCherry tag at the C-terminus of ParA-K20R (Figure 7A). Although reduced efficiency, we were able to construct the strain for expression of *parA(K20R-mcherry)* from the xylose promoter. Cells expressing the variant encoding ParA^K20R-mCherry^ displayed more severe defects in replication initiation and cell length compared to overexpression of *parA(WT)*. Notably, the expression of this ParA variant (ParA^K20R-mCherry^) also resulted in a significant increase of mini-cells (Figure 7B). Collectively, these data are supportive of a model where ParA itself has multiple roles that can be dramatically altered by expressing specific variants of ParA.

**Figure 7:**
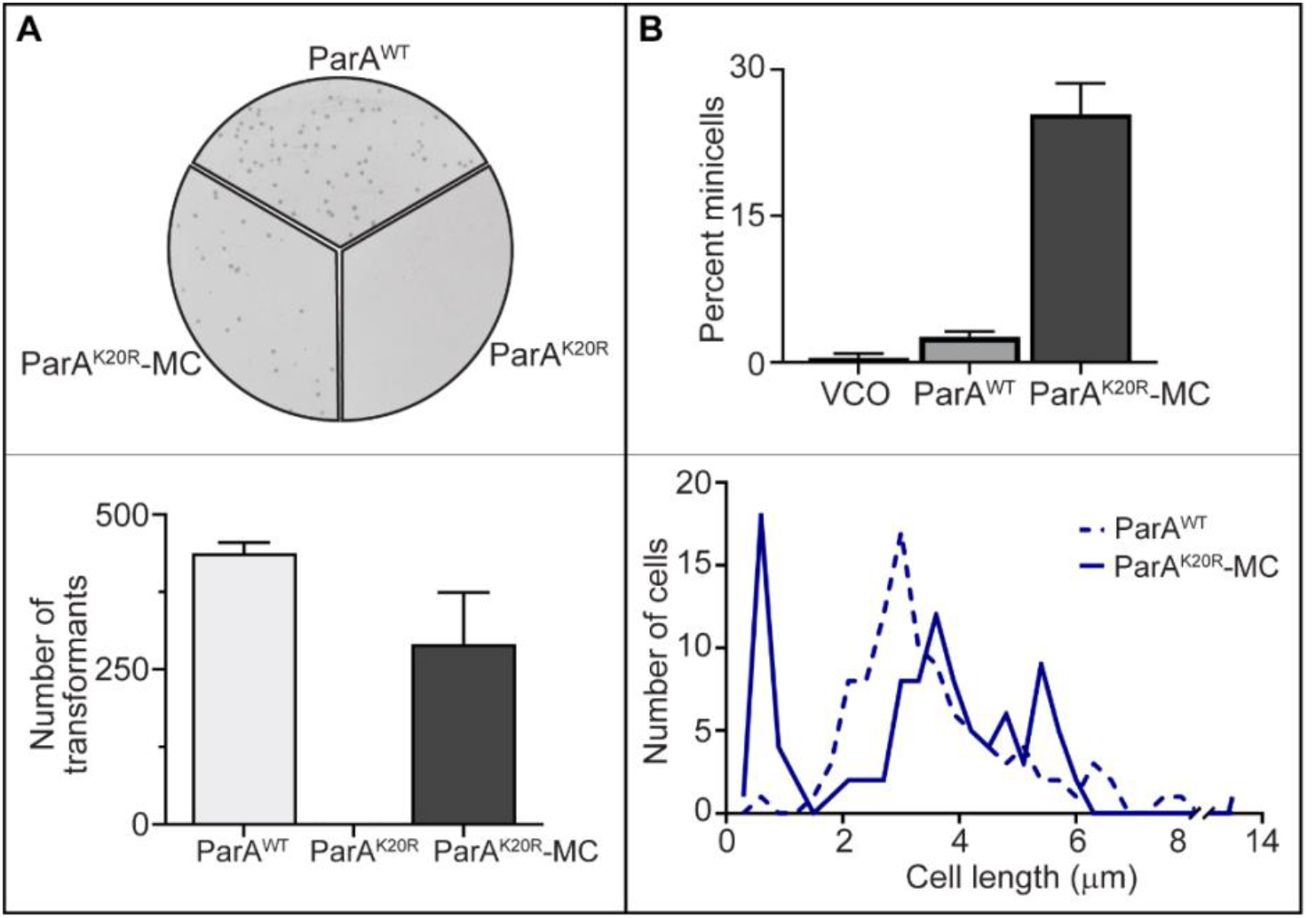
The monomer apo-ParA severely disrupts growth. (A) PYE/Kan agar plates with colonies resulted from the transformation of plasmids (100 ng) of *xylX::ParA*^*WT*^, *xylX::ParA*^*K20R*^ and *xylX::ParA*^*K20R*^*-MC* (mCherry tagged ParA) into *C. crescentus* CB15N *parB*:*cfp*-*parB* and the bar graph of quantification of transformants. No transformants were observed with *xylX::ParA*^*K20R*^ plasmid. Data represents the mean percent number ± SD of two independent experiments. (B) Bar graph of percent mini-cells (top) in cultures of CB15N *parB*:*cfp*-*parB* cells of *xylX::VCO* (empty vector control), *Par xylX::ParA*^*WT*^, and *xylX::ParA*^*K20R*^*-MC* grown in M2G + xylose (0.1 %) after 3h. Distribution of cell length of cells expressing ParA^WT^ and ParA^K20R^-MC for 3h in the presence of xylose (0.1 %). Micrographs taken at 3h were used to quantify the mini-cells and cell length. Quantification of mini-cells and cell length data are representative of three independent trials.

## DISCUSSION

Successful generation of a daughter cell requires the complex coordination of multiple events for proper progression of the cell cycle. The goal of this study was to identify the role of the partitioning protein ParA in coordinating other cell cycle events in *C. crescentus*. Our data revealed that ParA can promote the initiation of chromosome replication in a DnaA-dependent manner, but independent of ParA’s ability to promote chromosome segregation. ParA was unable to promote replication initiation when the predominant form of DnaA in the cell was in the deactivated form DnaA-ADP, suggesting that ParA is not involved in DnaA’s nucleotide activation step. Our data revealed that ParA’s impact on replication initiation does not change the regulation of CtrA levels over the cell cycle. Surprisingly, we discovered a potentially lethal effect from the apo-ParA monomer variant. Overall, our data provide new insights into the highly interconnected network that drives the forward progression of the bacterial cell cycle.

Our work adds *C. crescentus* with its dimorphic lifestyle to the number of bacterial species that use the partitioning system to fine tune complex developmental processes. For instance, in the stalk budding bacterium *Hyphomonas neptunium*, the par system is essential for regulating chromosome segregation in their unusual life cycle where replication and segregation of the chromosome are uncoupled [40]. In *B. subtilis*, the par system is essential during sporulation where Soj (ParA) prevents chromosome replication initiation in cells preparing to sporulate [13; 15]. In bacteria with multipartite genomes like *V. cholera*, the par system has been shown to coordinate the segregation of each chromosome with the onset of chromosome replication [41; 42]. Although the exact mechanisms seem to vary among bacterial species, the strategy of using the partitioning system to coordinate other cell cycle events involved in maintaining the integrity of the chromosome seems to be conserved in evolutionarily diverse species.

Our data suggest that aside from promoting chromosome replication initiation, ParA could also be involved with cell length regulation and/or cytokinesis. Cells with high levels of ParA^WT^ displayed increased cell length. Notably, cells expressing the dimer variant ParA^D44A^ displayed an even higher increase in cell length compared to ParA^WT^ while maintaining the same frequency of over-initiation of chromosome replication. These data suggest that different forms of ParA’s (i.e., monomers vs. dimers) may impact cell length independent of chromosome replication initiation. Additionally, cells expressing the variant apo-ParA-MC give off minicells at a higher frequency compared to the other variants. Collectively, these data posit a model where ParA can serve as a checkpoint in *Caulobacter’s* cell cycle. Various checkpoints have been identified that license the different cell cycle events in bacteria. CtrA is an example of such a checkpoint, which coordinates temporally and spatially cell development including chromosome replication initiation and cytokinesis [43; 44; 45]. Our work suggest that ParA is part of a new checkpoint in this complex system that is independent of CtrA that connects replication initiation, chromosome segregation, and cell length regulation (i.e., cytokinesis).

Our data revealed that altering the balance of ParA’s intermediates can be lethal to the cell. We were unable to construct a strain where the apo-ParA is expressed suggesting a potentially lethal effect on viability. Indeed, we found in the literature that whenever this variant of apo-ParA has been used in *Caulobacter*, it was fluorescently tagged [18; 22]. The striking effect from apo-ParA suggests at least two potential models. Firstly, apo-ParA could inhibit growth by binding ssDNA tightly preventing transcription and potentially chromosome replication. ParA-ADP has been shown *in vitro* to bind single stranded DNA (ssDNA) nonspecifically and potentially act as a transcriptional inhibitor [46]. The monomer apo-ParA may behave similarly to the monomer ParA-ADP. The second model is that apo-ParA-K20R impacts the cellular organization of the chromosome by interfering with ParB-SMC (Structural maintenance of chromosome complex) interactions. ParA-K20R has been shown to bind ParB more tightly than wildtype ParA [47]. Although the observed differences in the chromosomal organization were modest in the presence of ParA-K20R-YFP, the untagged ParA^K20R^ could have a more significant impact on chromosomal arrangement. In a recent preprint by Roberts, D. M. *et al*. (2021), the apo-ParA(Soj) was shown to control the positioning of SMC complexes impacting chromosomal arrangement during sporulation. These results are consistent with our findings that the apo-ParA may have specific functions in the cell. We are currently testing between these potential models in *C. crescentus*. More work is needed to determine the mechanisms of how different forms of ParA may influence specific events in the cell cycle.

## EXPERIMENTAL PROCEDURES

### Strains and Plasmids

All the strains, and plasmids used in this study are listed in Supplementary Table 1 and 2. Plasmids were constructed by cloning PCR amplified DNA fragments, into pXCHYC-2, pXCHYC-6, and pVCHYC-2 [48] vectors. *C. crescentus* wild-type CN15N (NA1000) genomic DNA was used as the PCR DNA substrate. Empty vector control plasmid was created by removing the mCherry tag sequence from pXCHYC-2 plasmid and self-ligating using Gibson reaction [49]. Plasmids were transformed into *E. coli* DH5a cells and grown in LB medium (Fisher Bioreagents, NJ) at 37°C with orbital shaking at 200 rpm. All plasmid constructs were verified by sequencing.

### Growth conditions

*C. crescentus* strains inoculated from freezer stocks were grown in M2G/PYE liquid media [50] at 30°C and 180 rpm. Liquid media was supplemented with 5 mg/mL Kanamycin (Kanamycin sulfate in water, IBI Scientific, NJ, USA), 0.2 mg/mL Chloramphenicol (in 100% ethanol) or 0.2 mg/mL Tetracycline (in 50 % ethanol). PYE plates contained 25 mg/ml Kanamycin or 1 mg/mL Chloramphenicol. Unless otherwise stated, *C. crescentus* cells were synchronized using the mini-synchrony protocol to isolate swarmer cells (first stage of the cell cycle). Briefly, cells grown in M2G (15 mL) to exponential phase (OD_600_ ∼0.3) were pelleted by spinning at 6000 rpm for 10 min at 4°C. Cell pellet was resuspended in 800 ml of 1 X M2 salts and then mixed with Percoll (900 mL, Sigma-Aldrich). The mixed solution in 2 mL microfuge tube was centrifuged at 11,000 rpm for 20 min at 4°C to separate the swarmer cells via density gradient separation. Bottom layer containing swarmer cells was carefully extracted out into a new tube, washed twice with 1 X M2 salts at 8000 rpm for 3 min at 4°C. Cells were resuspended in M2G as needed. As mentioned in the manuscript, when pre-induction of ParA/HdaA was required, the inducer 0.1% xylose / 250 mM Vanillate, (or amounts as noted, Sigma Aldrich) was added 1 h prior to synchrony.

### Growth analyses

Strains were inoculated from freezer stocks in M2G/PYE liquid media supplemented with Kan and incubated overnight at 30°C. Saturated cultures grown in M2G/PYE were diluted in fresh media (200 ml) to OD_600_ ∼ 0.01 in 96 well plates. Absorbance at OD_600_ was measured every hour in Biotek EPOCH-2 microplate reader at 30°C with orbital shaking at 180 rpm.

### Microscopy

Cells (∼ 2 mL) were spotted on agarose pads (1 % agarose in M_2_G). Agarose pads were supplemented with 0.1 % xylose during time lapse imaging. Phase-contrast fluorescence micrographs were obtained using Zeiss Axio Observer 2.1 inverted microscope with AxioCam 506 mono camera (objective: Plan-Apochromat 100×/1.40 Oil Ph3 M27 (WD=0.17 mm) and Zen lite software. Number of CFP-ParB foci per cell was manually counted (Cell Counter Plugin) and cell length was analyzed using ImageJ/FIJI with MicrobeJ software [51].

### Immunoblotting

*C. crescentus* swarmer cells were isolated using mini-synchrony protocol and the inducer was added as specified in the figure legends. OD_600_ of incubated cultures were normalized to ∼ 0.2 and the cell pellets resuspended in Cracking buffer (40 mL) were boiled at 80°C for 10 min and stored at −20°C for western blots. Proteins were separated on SDS-PAGE and transferred into the nitrocellulose membrane using iBlot2 (Invitrogen Dry Blotting System). The membrane was blocked with 1 X TBS (10 mM Tris-Cl, pH 8.0, 150 mM NaCl) with 5% non-fat milk and 0.1% Tween 20 (T) for 1h. Blot was probed with ∼1:10,000 diluted primary antibody (anti-Flag™ for ParA-M2, F7425 Sigma-Aldrich, and anti-DnaA for DnaA) overnight at 4°C. Membrane was washed three times with 1 X TBST and incubated for 1 h with 1:10,000 diluted secondary antibody (Anti-Rabbit IgG peroxidase, Sigma-Aldrich) at room temperature. Excess secondary antibody was removed by washing the membrane with 1 X TBST three times. Membrane was developed with SuperSignal™ West Pico PLUS Chemiluminescent Substrate (Thermo Scientific, IL) and imaged with ChemiDoc-MP imaging system (Bio-Rad Laboratories)

### Flow Cytometry Analysis

Swarmer cells were isolated via mini-synchrony (Tsai & Alley, 2001) and resuspended into M2G. At respective time points, aliquots were treated with a flow cytometry protocol previously described (Lesley & Shapiro, 2008) to analyze chromosome content. In summary, cell cultures were incubated with Rifampicin (15 μg/ml in 100% methanol) to block the re-initiation of DNA replication. The samples were incubated for 3 hours at 30°C. Afterward, they were fixed with 70% ethanol, and stored at 4°C for up to 24 hours. The fixed cells were then spun down at 4000 x g and resuspended gently in TMS buffer (10 mM Tris-HCl pH 7.2, 1.5 mM MgCl_2_, 150 mM NaCl) and stained with 10 μM Vybrant DyeCycle Orange (Invitrogen). The fixed samples were analyzed on a BD LSR II Flow cytometer. Flow cytometry data were analyzed using BD FACSDiva software (BD Biosciences).

## ACKNOWLEDGEMENTS

We are appreciative to the generosity from Dr. Kristina Jonas for sharing her strain KJ300 and from Dr. Justin Collier for sharing her HdaA antibodies. We thank Willow Boecklen for his assistance with the initial data analyses and Israel Ponce for the construction of Figure 1. The work reported in this publication was supported by the National Institute of General Medical Sciences of the National Institutes of Health under Award Number R01GM133833. S.G. P-R. was supported R01GM133833-Supplement and R. A. E. was supported by the National Institute of Health – Maximizing Access to Research Careers (MARC) (GM07667-39).

