## Supplemental Figures for "ParA and its functions that go beyond chromosome segregation in *Caulobacter crescentus*"

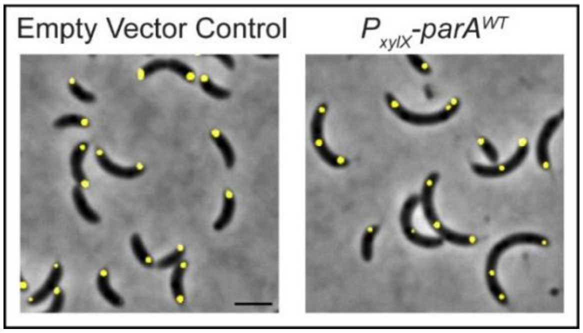


**Figure S1: Over-initiation of chromosome replication in *cc0006::(tetO)n (gentr), Pvan:tetR-eyfp C. crescentus* cells expressing ParA$^{WT}$.** *C. crescentus cc0006::(tetO)n (gentr), Pvan:tetR-eyfp* swarmer cells having empty vector or *xylX::parA$^{WT}$* in M2G (2 ml) supplemented with 0.1 % xylose were incubated at 30°C in a roller-shaker. Phase-contrast fluorescence micrographs of cells expressing ParA$^{WT}$ obtained at 4h shows more than 2 TetR-YFP yellow foci corresponding to *oris*/over-initiation compared to empty vector control with 2 TetR-YFP foci. Micrograph scale bar - 2μm. Cells shown are representative of three independent experiments.

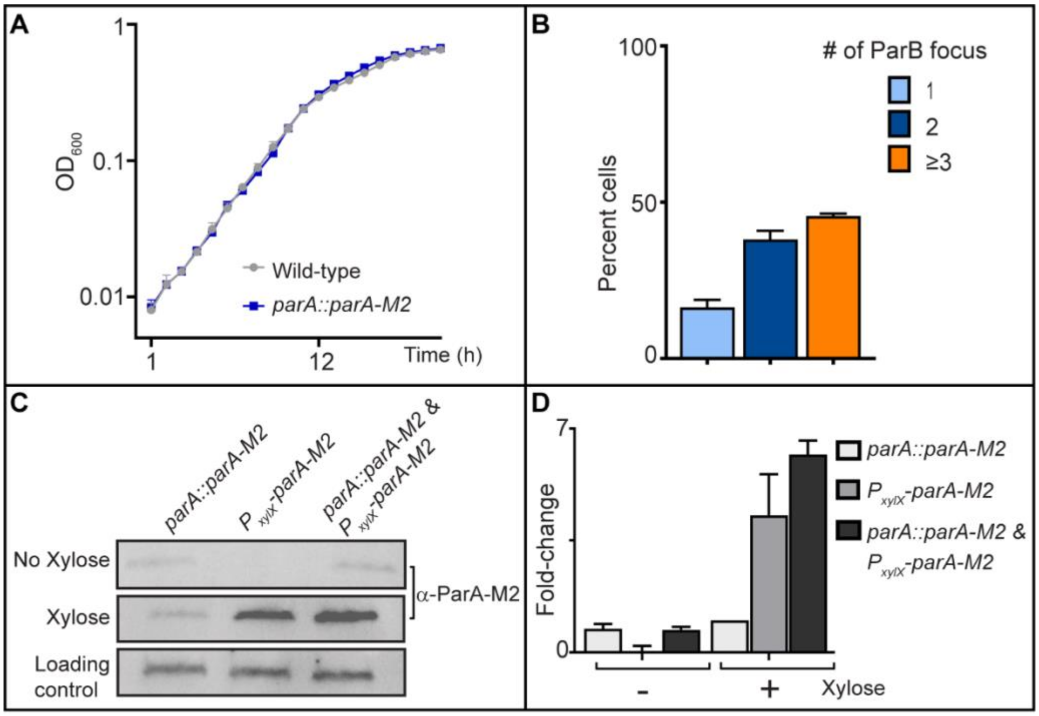


**Figure S2: Characterization of ParA-M2 and overexpression levels.** (A) Growth curves of *C. cresentus* CB15N *parB::cfp-parB* (wild-type) and *parA::parA-M2* cells in M2G media. (B) Quantification plot of CB15N, *xylX::parA-M2* cells expressing $ParA^{WT}$-M2 in the presence of xylose (0.1 %) for 3 h. (C) Western blots of ParA-M2 levels expressed under native promoter (no xylose) and in the prescence of xylose after 3 h. (D) Bar graph of quantfication plots of western blots (C). A and C - Data represent independent three experiments. B and C data are from three independent experiments with error bars of mean ± standard deviation (SD).

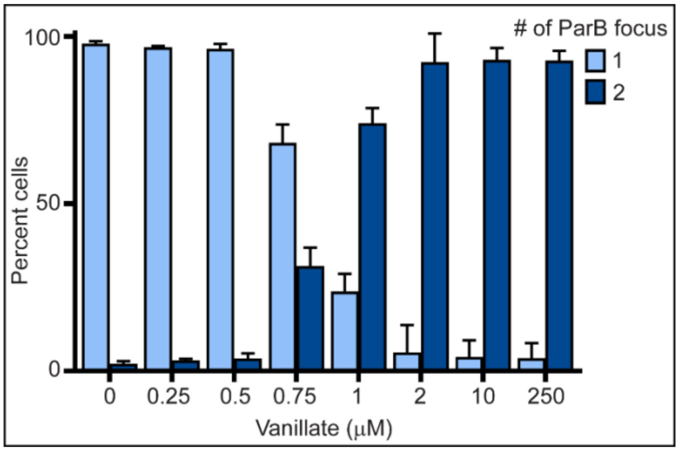


**Figure S3: Quantification of frequencies of replication initiation in cells with titrated *dnaA* induction.** Bar graph of percent of replication initiation of *C. crescentus* CB15N, *parB*::*cfp-parB, ΔvanA, dnaA*::Ω, *vanA*::*dnaA* cells under increasing amounts of vanillate (0 -250 μM) to induce DnaA. Swamer cells in M2G (2 ml) were incubated at 30°C in a roller-shaker for 3 h to deplete DnaA and added increasing amounts of vanillate (0, 0.25, 0.5, 0.75, 1, 2, 10 and 250 μM) to induce DnaA expression. Micrographs were obtained after 1 h incubation and the percent cells CFP-ParB was quantified. Data are from three independent experiments with error bars of mean ± standard deviation (SD).

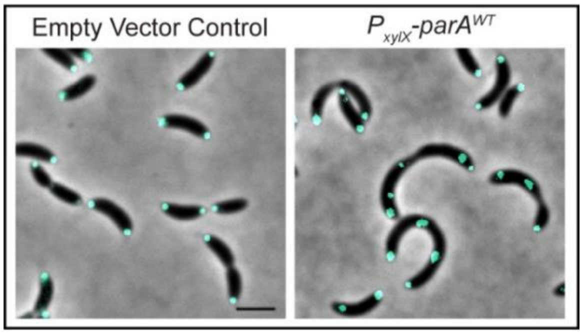


**Figure S4: *C. crescentus* grown in rich media (PYE) expressing ParA$^{WT}$ promotes over-initiation of chromosome replication in *C. crescentus*.** Overnight cultures of *C. crescentus* CB15N*, parB::CFP-parB* cells having empty vector or *xylX::parA$^{WT}$* grown to exponential phase in PYE rich media (5 ml) were used to set $OD_{600}$ ~ 0.1 in fresh PYE rich media (2 ml) supplemented with xylose (0.1 %). Phase-contrast fluorescence micrographs of cultures incubated at 30°C in a roller-shaker were obtained at 3h. Over-initiation of replication was observed in cells expressing ParA$^{WT}$ (more than 2 CFP-ParB foci/*oris*) compared to empty vector control. Micrograph scale bar - 2μm. Cells shown are a representative of three independent experiments.

**Table 1:** List of strains used in this study

| **Name** | **Relevant genotype or description** | **Reference** |
|---|---|---|
| ***C. crescentus* strains** | | |
| | CB15N (NA1000) | [1] |
| PM109 | CB15N, *parB*::*cfp-parB,* Δ*vanA*, *dnaA*::Ω (strp$^R$/spec$^R$), *vanA*::*dnaA* | [2] |
| PM117 | CB15N, *parB*::*cfp-parB, xylX::parA*$^{K20R}$*-mCherry* (tet$^R$) | This study |
| PM258 | CB15N, *parA::parA-M2* | This study |
| | CB15N, *parB*::*cfp-parB* | [2] |
| PM502 | CB15N, *parB*::*cfp-parB, xylX::parA-mCherry* (kan$^R$) | This study |
| PM541 | CB15N, *parB*::*cfp-parB, xylX::parA*$^{WT}$ (kan$^R$)*, parB::cfp-parB* | This study |
| PM542 | CB15N, *parB*::*cfp-parB,* Δ*vanA*, *dnaA*::Ω (strp$^R$/spec$^R$), *vanA*::*dnaA, xylX::parA*$^{WT}$ (kan$^R$) | This study |
| PM550 | CB15N, *parB*::*cfp-parB, xylX::parA*$^{D44A}$ (kan$^R$) | This study |
| PM566 | CB15N, *parB*::*cfp-parB, xylX::empty-vector* (kan$^R$) | This study |
| PM607 | CB15N, *parB*::*cfp-parB,* Δ*vanA*, *vanA*::hda*A* (chlor$^R$)*, xylX::parA*$^{WT}$ (kan$^R$) | This study |
| PM651 | CB15N, *parB*::*cfp-parB, xylX::parA*$^{R195E}$ (kan$^R$) | This study |
| PM656 | CB15N, *parB*::*cfp-parB, xylX::parA*$^{R195A}$ (kan$^R$) | This study |
| KJ300 | CB15N *cc0006::(tetO)*$_n$ (gent$^R$ )+P*xylX::lacI-ecfp, tetR-eyfp* (spec$^R$) (gent$^R$ , spec$^R$) | [3] |
| PM714 | CB15N, *xylX::parA-M2* (kan$^R$) | This study |
| PM718 | CB15N *parA::parA-M2, xylX::parA-M2* (kan$^R$) | This study |
| PM750 | CB15N, *parB*::*cfp-parB, ΔparA*, *xylX::parA-M2* (chlor$^R$) | This study |
| PM767 | CB15N, *xylX::parA*$^{G16V}$ (kan$^R$)*, parB::cfp-parB* | This study |
| PM771 | CB15N, *parB*::*cfp-parB,* Δ*vanA*, *dnaA*::Ω (strp$^R$/spec$^R$), *vanA*::*dnaA, xylX::empty-vector* (kan$^R$) | This study |

Table 2: List of plasmids used in this study

| **Plasmid name** | **Description** | **Reference** |
|---|---|---|
| pXCHYC-2 | Integrating constructs encoding C-terminal mCherry fusions under the control of native $P_{xylX}$ (kan$^R$) | [4] |
| pVCHYC-6 | Integrating constructs encoding C-terminal mCherry fusions under the control of native $P_{vanA}$ (chlor$^R$) | [4] |
| pXCHYC-5 | Integrating constructs encoding C-terminal mCherry fusions under the control of native $P_{xylX}$ (tet$^R$) | [4] |
| pDNA91 | *parA-mCherry* cloned into pXCHYC-2 | This study |
| PM116 | *parA (K20R)* cloned into pXCHYC-5 | [5] |
| pDNA245 | *parA* cloned into pXCHYC-2 | This study |
| pDNA255 | pNPTS138 derivative to replace *parA* by *parA-M2* under the native $P_{parA}$ | This study |
| pDNA257 | *parA (D44A)* cloned into pXCHYC-2 | This study |
| pDNA264 | *mCherry* tag excised from pXCHYC-2 | This study |
| pDNA274 | *hdaA* cloned into pVCHYC-6 | This study |
| pDNA309 | *parA (R195E)* cloned into pXCHYC-2 | This study |
| pDNA310 | *parA (R195A)* cloned into pXCHYC-2 | This study |

| | | |
|---|---|---|
| pDNA315 | *parA-M2* cloned into pXCHYC-2 | This study |
| pDNA321 | pNPTS138 derivative to delete native *parA* while keeping the downstream *cfp-parB* | This study |
| pDNA323 | *parA-M2* cloned into pXCHYC-6 | This study |
| pDNA329 | *parA (G16V)* cloned into pXCHYC-2 | This study |
